# Cumulating MS Signal enables polyclonal antibody analysis

**DOI:** 10.1101/2025.03.31.645874

**Authors:** Carlos Gueto-Tettay, Joel Ströbaek, Di Tang, Alejandro Gomez Toledo, Yasaman Karami, Hammed Khakzad, Johan Malmström, Lars Malmström

## Abstract

Unraveling the complexities of protein systems via Mass Spectrometry (MS), particularly polyclonal antibodies, demands innovative analytical strategies. Here, we introduce the cumulative MS score (cMS), a novel mathematical framework that transcends traditional spectrum-matching, integrating MS evidence across multiple sample injections to achieve robust *de novo* peptide sequencing annotation. This approach, shifting from isolated spectrum analysis to a holistic MS signal-based methodology, was rigorously evaluated and validated across diverse sample types and experimental conditions. We applied this framework to characterize a complex polyclonal antibody mixture of *Streptococcus pyogenes* M1 protein binders derived from intravenous immunoglobulin (IVIG), revealing predominant variable heavy (VH) and light (VL) chain subgroups consistent with established genetic studies. Furthermore, we successfully identified conserved complementarity-determining region (CDR) features and predicted stable antibody-antigen interactions through molecular dynamics simulations, demonstrating the method’s potential for dissecting intricate antibody responses. This work establishes a powerful alternative to conventional tandem mass spectrometry MS/MS data analysis, enabling deeper insights into protein systems and paving the way for targeted therapeutic development.

## Introduction

Tandem mass Spectrometry (MS) has revolutionized molecular biology by providing unprecedented sensitivity and precision in analyzing complex biological systems^1–4^. This powerful analytical technique enables researchers to characterize proteins, peptides, and other biomolecules with extraordinary detail^5^, detecting minimal quantities of molecular components that would be undetectable through traditional analytical methods^6,7^. From proteomics^8^ and metabolomics to clinical diagnostics^4,9^ and drug development, MS has become an indispensable tool for investigating the intricate molecular landscapes of living systems^1,2,9,10^. The inherent challenge in many biological research scenarios lies in the lack of comprehensive reference databases, particularly when studying unique or poorly characterized molecular systems. *De novo* MS peptide sequencing emerged as a transformative approach to address this limitation, offering a method to directly interpret proteomics mass spectra and determine peptide sequences without relying on existing protein databases^11–13^. This technique becomes especially crucial when investigating complex proteins like antibodies, where molecular diversity and genetic variability create significant computational challenges^14–17^.

Antibodies (immunoglobulins) are crucial adaptive immune system components that pose unique analytical challenges due to their inherent complexities^18^. They achieve high specificity in binding their cognate target (antigen) through the variable fragment (Fv) region, a product of intricate genomic processes. The Fv region, encompassing the variable heavy (VH) and variable light (VL) domains. It also features four framework regions (FR1-4) that scaffold the three complementarity-determining regions (CDRs), which directly mediate antibody-antigen interaction. Understanding these interfaces, known as the paratope (antibody) and epitope (antigen), is essential for developing targeted immunotherapies. Current experimental methods struggle to effectively bridge the gap between antibody genetic elements, protein assembly, maturation, and final target epitope specificity. Next-generation sequencing methods, such as B-cell receptor sequencing (BCR-seq), face limitations in resolving heterodimer assembly without costly or laborious single-cell techniques and may not fully represent circulating antibody populations^19,20^. Furthermore, the sparsity of structural data for antibody-antigen complexes hinders protein-protein interaction prediction^21,22^.

In this context, *de novo* MS/MS sequencing presents a promising strategy to directly connect antibody sequencing with structure-based interface analysis. Addressing current throughput limitations could significantly expand available structural epitope-paratope information, paving the way for future deep learning-driven advancements in antibody research. Since 2017^23^, the field has witnessed a remarkable surge in deep-learning-based *de novo* MS peptide sequencing tools, each promising to enhance our ability to interpret complex mass spectrometric data^11–16,20,23–37^. However, the prevailing analytical approach has remained fundamentally limited, typically analyzing each sample and spectrum in isolation. This approach fails to leverage the rich, redundant information present across multiple experimental injections and sample preparations. For better understanding, let’s consider a protein with a known sequence submitted to a multi-enzyme digestion protocol. Every protease creates its own set of peptides, showing off protein’s many variations and hidden details, like post-translational modifications. Furthermore, a proteomics analysis surely reveals a network of overlapping peptides that comprehensively map the protein’s primary structure. From the MS perspective, we now have a rich fragment ion signal for all sequence positions of the target protein. Existing *de novo* sequencing methodologies predominantly analyze individual mass spectra in isolation, systematically disregarding the protein-level contextual information. This reductionist analytical approach intrinsically limits the comprehensive interpretation of molecular diversity, manifesting as substantial variability and inconsistency across different *de novo* MS peptide sequencing tools. In other words, the mass spectra missing fragment ions events can lead to the creation of multiple conflicting candidate peptides that can be explained by the same MS evidence, called here ranking ambiguity cases. The latter translate into a common scenario, where different *de novo* MS peptide sequencing tools reporting different sequences as their top1 scored candidates. The prevailing methodological constraint emerges from the absence of an integrative framework capable of synthesizing spectral information within the broader protein sequence context. This limitation is further compounded by the need to leverage the diverse features offered by various MS-based *de novo* sequencing tools. Specifically, methods must account for the varying capabilities of these tools, including their ability to consider specific post-translational modifications and their overall *de novo* sequencing accuracy, particularly when dealing with the substantial datasets required for robust model training. Consequently, the current analytical paradigm fundamentally constrains our ability to extract nuanced molecular insights from complex proteomic datasets.

We propose a novel method that transcends current *de novo* MS/MS sequencing limitations by introducing the cumulative MS score (cMS), representing a paradigm shift in MS data analysis. By integrating fragment ion information across all injected MS samples, we propose a comprehensive approach that significantly improves peptide annotation processes. This method enhances the reliability of peptide identification, and by extension, primary structure assignment. It also provides a flexible framework for incorporating future developments in computational analysis. Specifically, our approach leverages cumulative and redundant fragment ions information to extract critical molecular features from complex biological systems (Figure 1). To rigorously validate our approach, we conducted comprehensive experiments using commercially available antibodies with well-characterized sequences, including Herceptin, Xolair, Prolia, and Avastin. These validation studies involved systematic analysis across multiple experimental conditions, including different proteases, digestion times, and MS acquisition modes. By leveraging the cumulative and redundant fragment ion information acquired across all injected samples, our method significantly improves the annotation process, demonstrating superior performance compared to traditional isolated analysis techniques at both the scan and peptide levels. We then demonstrate the power of this methodology through a comprehensive analysis of a polyclonal antibody population, targeting the clinically relevant pathogen *Streptococcus pyogenes*^38–41^, showcasing how our cumulative approach can unveil molecular insights that traditional isolated analysis methods would miss. This included position-specific CDRH3 features that *in silico* demonstrated a higher affinity for distinct *S. pyogenes* M1 protein epitopes, which could have implication for designing targeted immunotherapeutics. By moving beyond individual spectrum interpretation and string-based analysis to a MS signal data-driven approach, we open new avenues for understanding molecular diversity and interaction dynamics in intricate biological systems. cMS addresses current limitations for *de novo* MS data analysis, while providing a universal framework applicable across diverse research domains, from immunology and protein characterization to emerging fields of molecular biology.

**Figure 1.**
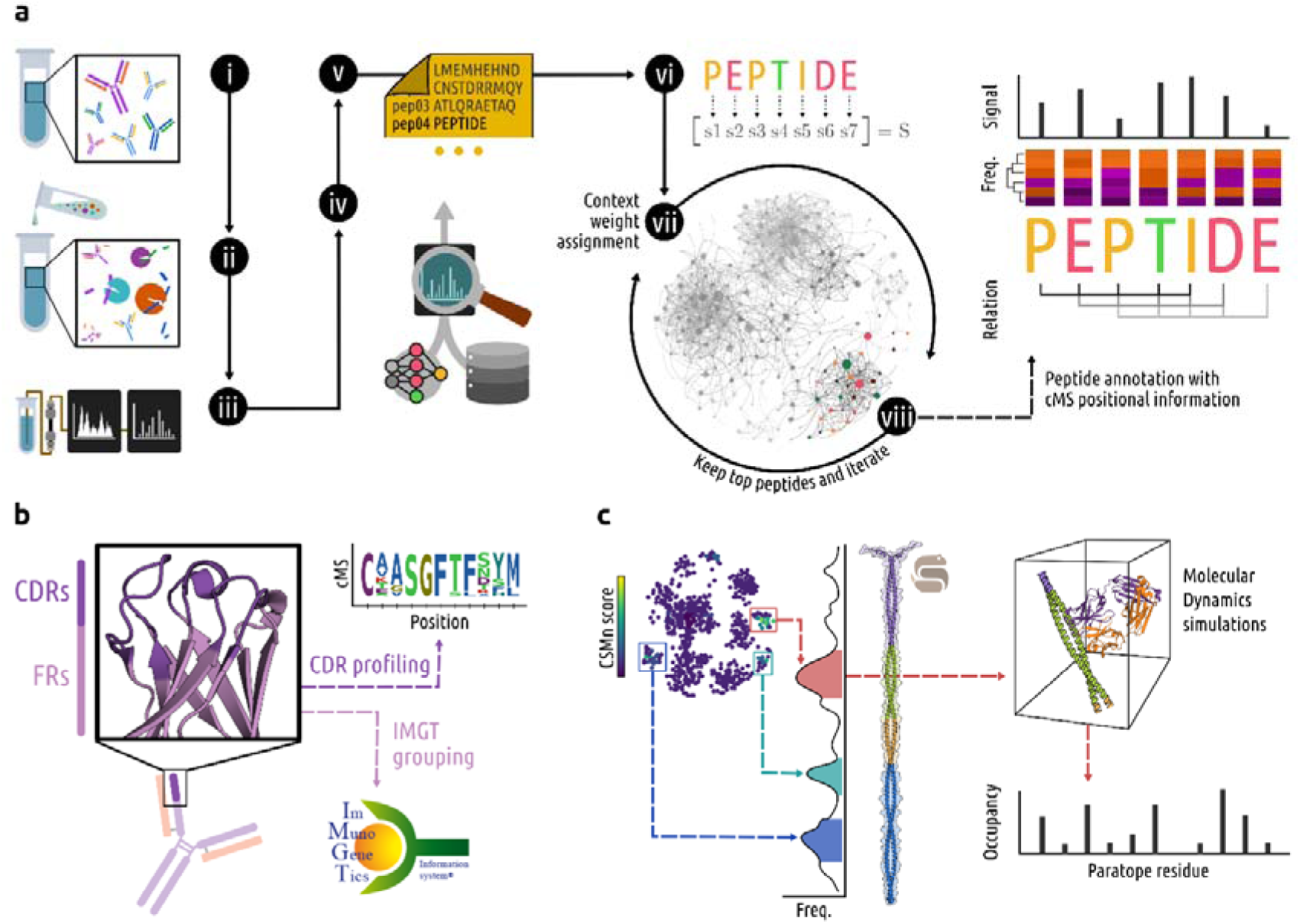
General cMS annotation and applications workflows. **a)** protein samples (i) are submitted to the multi-enzyme digestion protocol (ii). Peptides are separated and detected via LC-MS/MS(iii). We created an expanded candidate list for each spectrum using several *de novo* and database search engines (iv and v). For each candidate peptide, we initially calculate its isolated score. Then, the context weight component is obtained by integrating the MS signal from all available candidates. Finally, we calculated the weighted scoring function for all candidates. We iteratively kept the top N high-scored until only one candidate peptide per spectrum remains (vii and viii). **b)** The resulting cumulated MS vectors are used to study and analyze the variable domain features of the polyclonal antibody population, specifically to identify the most abundant IMGT group and subgroups and the CDRs feature space distributions. **c)** We coupled the data with protein-protein docking and MD simulations to explore the CDRH3 space for a deeper insight into the paratope-epitope interface.

## Results and discussion

We set up a series of experiments to assess the generalizability and robustness of a novel cumulative MS evidence-based method. Given MS/MS samples coming from the multi-enzyme digestion of a target protein with a known sequence, the goal of the experiments was to evaluate the scoring method’s ability to identify the true-positive peptides from a set of true-negative decoy peptides. Several spectrum match scoring functions were tested with conventional methods (the isolated way) and a novel context-weight method called cumulative MS (cMS; see *Computational details*).

### cMS produce higher scan and peptide recall

In the first experiment, we showcased the applications of our method using the commercial antibody Herceptin. We retrieved our previously reported MS/MS data of multi-enzyme digested Herceptin^42^, generated with diverse fragmentation settings, from the ProteomeXchange Consortium (PRIDE, PXD037803). We then created candidate peptides through a spectrum-matching database search approach. All 55 samples were submitted to comprehensive, nonspecific-termini, searches against a database created from the combination of the entries of the target protein chain sequences and five species proteomes (see *Computational details* section). To assess the quality of the spectrum-matching peptides we devised a score (called S4b), for which we considered the sum of the amount of observed fragment ions and their intensity normalized by their peptide length. We repeated the experiment using candidate inclusion criteria with a minimum amount of found fragment ions (ranging from1 to 6).

Figure 2 shows the efficacy of our method for improving the spectrum-matching process. Figure 2a shows that the context-weighted scoring system identified more scans than the traditional method for all values of the experimental inclusion criteria, with differences ranging from 1.9–3.1 times. These results also showed an expected decrease in the target spectra as we increased the minimum requirements for inclusion. The cMS approach thereby demonstrated a high capacity for correctly annotating the dynamically available target spectra space. In contrast, the Isolated method flat line suggests that the primary source of contribution was easily fragmented peptides with rich fragment ion information, annotating up to 156,375 matching spectra with the most permissive inclusion criteria (Figure 2a). It is worth pointing out that this number was always surpassed by the cMS method, even for the most restrictive tested inclusion criteria.

**Figure 2.**
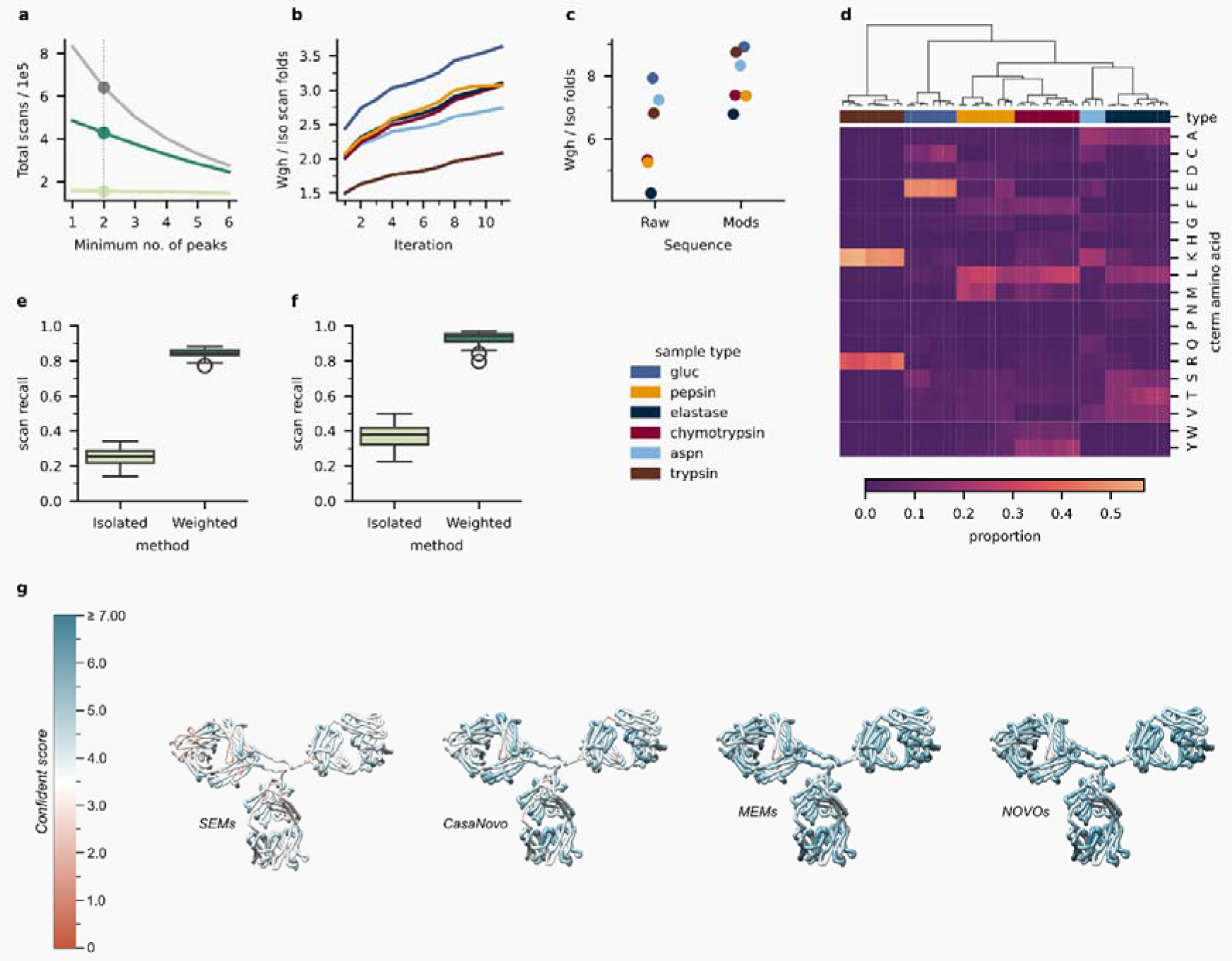
Isolated (Iso) vs weighted (Wgh) scoring system spectrum-annotation results from the Herceptin test set. **a)** The effect of the minimum number of matching fragment ion peaks (n_peaks) as inclusion criteria for the number of annotated scans. The grey line represents the total number of scans bearing a Herceptin-matching peptide. Regarding the line color scheme, the dark-green, light-green lines are the number of scans obtained from using the weighted and isolated scoring systems, respectively; Panels **a** - **d** explore details from taking n_peaks >= 2 results **b)** Sample-type Wgh/Iso annotated scans proportion in every iteration; **c)** Wgh/Iso proportion of detected raw (unmodified) and modified peptide sequences; **d)** C-terminus amino acid distribution for all sample types. Scan recall values from testing different S1, S4, S4b, and S4w scoring functions on **e)** a combination of database and *de novo* results data and **f)** *de novo* results only; **g)** the effect of integrating results from different deep-learning models (SEMs and MEMs) and tools (DeepNovoV2 and CasaNovo) on the protein confident scores. *NOVOs* combine all the aforementioned *de novo* peptide results into one expanded candidate peptide list.

A benefit of our proposed method is entirely suppressing the otherwise commonly observed ranking ambiguity cases, as changes in amino acid sequence affect the overall context weight value (see *Cumulative MS scoring system* section). This was not the case for the Isolated method, where 4.6–6.2% of the identified scans contained one or more additional candidate peptides with the same S4b score. Other benefits include improvements for annotating scans, uncovering peptide diversity, clustering, and identifying cleavage rules for all enzyme sample types. To elaborate on these points, let us consider the cases where the minimum number of fragment ions found was two. Figure 2b shows how the iterative scan assignment strategy boosted the matching candidate peptide identifications, compared to the Isolated method. This was accomplished while effectively removing decoys in an unbiased data-driven manner, and resulted in a final increase in sequence recovery of 2–3.8 folds compared to the standard scoring methodology. A downstream consequence of this was the 4–8 times expansion of the Herceptin matching peptide space, as shown in Figure 2c, which was further reinforced by the augmentation of the results when considering all peptide forms. Uncovering peptide diversity is of great value in the *de novo* MS protein sequencing field, where finding overlapping peptides is crucial for confident primary structure rebuilding and validation. Figure 2d highlights the ability of the cMS to cluster sample types, showing overall similar intra-sample type behavior. Moreover, C-terminus amino acid distribution data suggests that our approach enables the unbiased identification of the protease cleavage sites. The most noticeable results correspond to enzymes with more strict cleavage rules, like trypsin and gluc, where positive (K and R) and negative (E) charged amino acids were found in an elevated proportion. Other enzymes present multiple amino acids, such as elastase (A, L, S, T, V) and chymotrypsin (F, H, L, M, W, Y). Clustering sample types by the N-terminus amino acid proportions leads to similar grouping patterns, accentuating D amino acid for the aspn sample type (*Supplementary information*).

### Integrating more de novo MS sequencing tools led to higher confident scores

One of the most important features of our approach is the ability to combine results from different MS peptide sources. This is especially relevant given the recent growth in deep learning-based *de novo* MS peptide tools and models. For systems that include highly conserved amino acid sequences, such as monoclonal antibodies, it is also possible to contribute partial information from conventional database searches. To showcase this, we challenged our proposed method by expanding the candidate peptide lists by appending the *de novo* MS tools DeepNovoV2 and CasaNovo results. For DeepNovoV2, we used several previously trained Single-Enzyme (SEM) and Multi-Enzyme (MEM) Models, taking the top 200-scored candidates for each spectrum. In addition, the top 20 results were generated with the default CasaNovo model (see *Peptide candidates list generation* section). Here, we measured the scan recall or proportion of spectra where the best-ranked candidate peptide belonged to the target protein. Figure 2e shows that the increased peptide space did not affect the cMS approach for identifying matching peptides. As expected, the Isolated method identified fewer MS spectra and more ambiguous cases than the database-only setup. The latter results are explained when considering that many spectra contain missing fragment ions. Unlike database searches, where candidate peptides are restricted to a designated proteome sequences space, *de novo* MS peptide search tools introduce ambiguity by proposing amino acid sequences that can be explained by the same pool of available fragment ions. Repeating the experiment with only *de novo* data led to the same conclusions regarding scan annotations and peptide diversity, as shown in Figure 2f. Moreover, Figure 2g displays how we increased the Herceptin primary structure evidence by progressively integrating deep learning-based *de novo* MS peptide sources. We extended this experiment to other commercial antibodies, under a variety of experimental and computational settings, yielding similar conclusions to those found in Figure 2. It is also worth mentioning that utilizing the methodology established in NOVOs, we successfully *de novo* sequenced an antibody targeting the *S. pyogenes* antigen, Streptolysin O (SLO), which was subsequently used for epitope mapping and structural analysis, as detailed in Tang *et. al., 2024*^10^.

### cMS works across many computational and experimental settings

We continued to challenge our MS spectrum-matching scoring framework with 98,304 new *in silico* experiments (Figure 3). Each one consisted of an assortment of four parameters: MS acquisition mode, digestion times combinations, scoring scheme method, and scoring function. The commercially available antibody Avastin was subjected to digestion with six different proteases, with samples collected at three distinct time points. A pooled sample of all time points was also generated. We also introduced alpha-lytic in substitution of aspn using the same sample preparation protocol. The MS data was acquired using data-dependent acquisition (DDA) and data-independent acquisition (DIA) modes (see *Material and methods section*). The annotation spectra process was done with four scoring functions: S1, S4, S5, and E-score (see *Computational details* section). Like previous experiments, we assessed the ability of the cMS score to annotate the spectra of interest correctly (Figure 3). Three things are worth noting from this round of experiments. Firstly, scan recall values ranging from 0.90–0.99 reinforce the idea of the cumulative MS approach as a superior method for *de novo* MS sequencing peptide annotation compared to the isolated scoring method (Figure 3a). Secondly, it also shows that the method can work by accumulating signals from different MS spectrum attributes as reflected in the scoring functions tested, e.g., counting the presence of fragment ions (S5) and intensity (S4) and weighting the signal from the m/z deviation (S1). Lastly, it indicates that performing *de novo* MS peptide sequencing is feasible directly from MS data acquired with DIA mode or combined with DDA data (here called DXA). Compared to the DDA acquisition method results, the DIA and DXA data revealed a boost in the target protein MS evidence, i.e., 4.1 and 1.8 folds at the spectra annotation (Figure 3b) and matching peptide identification (Figure 3c) levels, respectively. Overall, the method demonstrated robustness across a wide range of experimental and computational setups, evidenced by consistently high scan recall and enhanced target protein MS evidence despite variations in acquisition mode, digestion protocols, and scoring functions.

**Figure 3.**
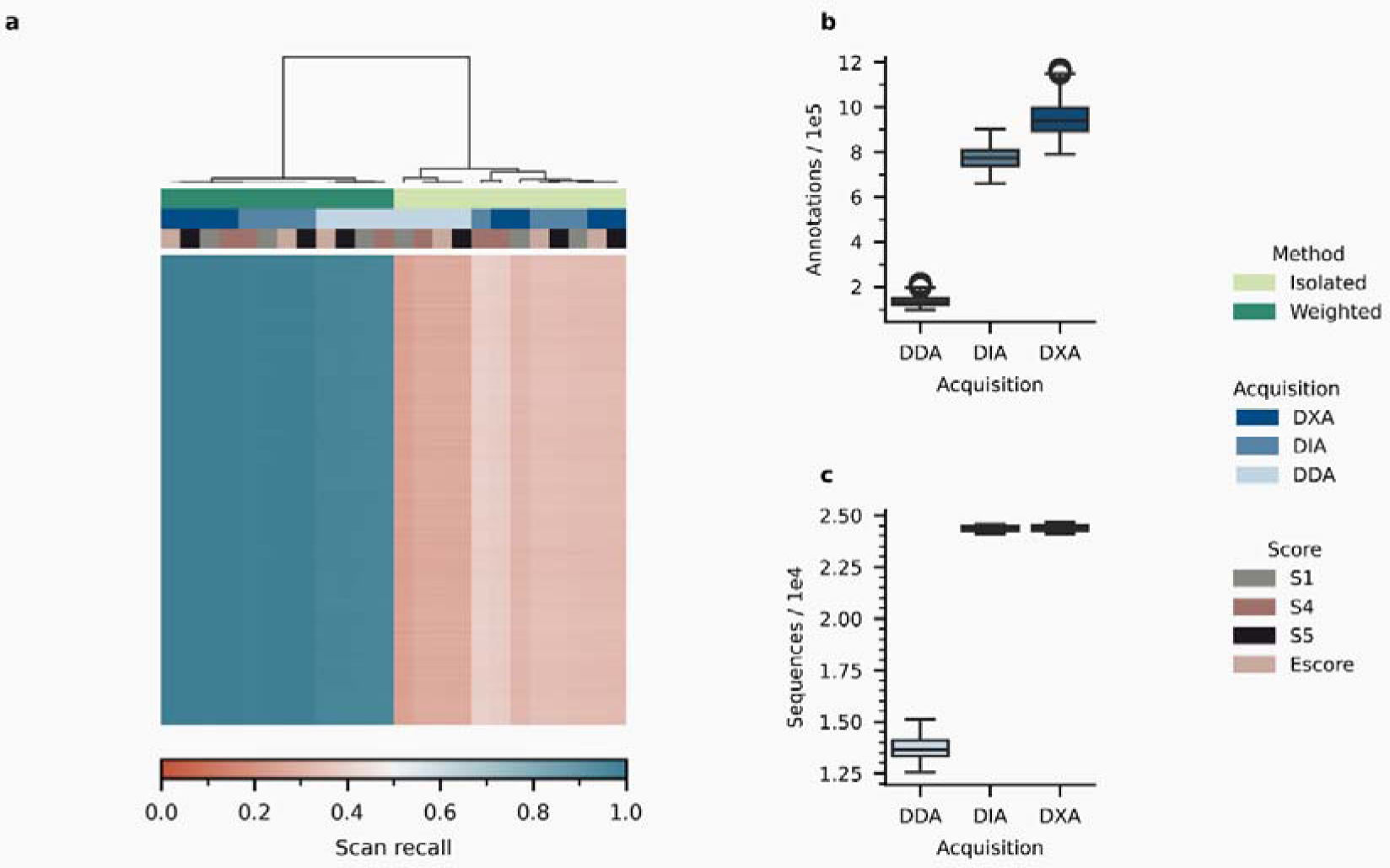
Multi-parameter Avastin dataset. **a)** Scan recall for 98304 in silico experiments from selecting a combination of different acquisition methods (DDA, DIA, DXA), scoring system method (Isolated and Weighted), scoring functions (S1, S4, S5, and E-score) on a combination of sample-type and digestion times. e.g., We can combinate the following MS samples: trypsin-4h, chymotrypsin-9h, pepsin-30min, elastase-overnight, and alpha-lytic-pooled samples. We then calculate the scan recall from the *de novo* MS peptide sequencing results by combining the remaining parameters. In addition, we also compared the DDA, DIA, and DXA methods at a **b)** scan-level and **c)** sequence-level.

### Positional cMS vectors unlock antibody population analysis

After comprehensively assessing our newly developed *de novo* MS peptide sequencing annotation method, we hypothesized that a cumulative MS signal analysis would garner valuable insights into more complex protein systems. We moved from a string-based to an MS signal-based analysis to better characterize the molecular system, as higher cMS values denote molecular species with higher experimental MS evidence. To test this hypothesis, we chose to characterize antibodies against *S. pyogenes* (also called Group A Streptococcus or GAS). GAS is a human-specific bacterium that challenges the human immune system throughout their lifetime, primarily evident in adolescence through pharyngitis. This means that the majority of healthy adults carry antibodies that target GAS antigens. We narrowed our analysis further by focusing on the Streptococcal M protein, which is a cell wall-anchored dimeric coiled-coil, and a well-characterized antigen critical for GAS pathogenicity ^43^, with the ability to form interaction networks with plasma proteins to lower immunogenicity ^39,43,44^. These include interfaces with antibodies in different binding modes ^45–47^ along its repetitive sequence of residues, with some domains being the focus of ongoing vaccination attempts^38^. We prepared a complex polyclonal mixture by pulling down GAS M1 protein binders from Intravenous Immunoglobulin (IVIG). IVIG is a concentration of pooled immunoglobulins derived from 1000–100,000 healthy donors^48^, which should contain antibodies targeting GAS and, by extension, the M1 protein^49,50^. To further limit our analysis to immunogenically relevant interaction, we focused our efforts on the antibody Fv region (Figure 1b).

Our first objective was to find out what type of Fvs were detectable (Figure 4). To this end, the IMGT^51^ (international ImMunoGeneTics information system®) annotation scheme was used to classify the identified groups and subgroups for both heavy (VH) and light (VL) chain Fv domains. We specifically used the FR1–3 conserved regions to represent the identified clusters and fragmented every identified peptide into 7-mers and element-wise pooled the fragment ion information across all MS samples. Then, we mapped the 7-mers onto the frameworks in a human IgG database formed by 669,365,000 sequences (272,154,774 heavy, and 397,210,226 light chains) retrieved from the Observed Antibody Space (OAS) resource^52^. Figure 4a displays the normalized positional signal for both chains. VH results show higher overall evidence for the IGHV3 group, specifically the IGHV3-23 and IGHV3-30 subgroups. Similarly, VL results show a strong signal from IGKV1 and IGKV3 groups, particularly the IGKV3-20 and IGKV1-39 subgroups (*Supplementary information*). These findings are consistent with reported proportions of group/subgroup distributions in healthy individuals^53,54^, where the IGHV3 group in particular has a low association with disease^55^, and IGHV3-23 is over-represented in naive B-cells^56^. This also reflects the state of the experimental setup, which utilized antibodies sourced from healthy adults.

**Figure 4.**
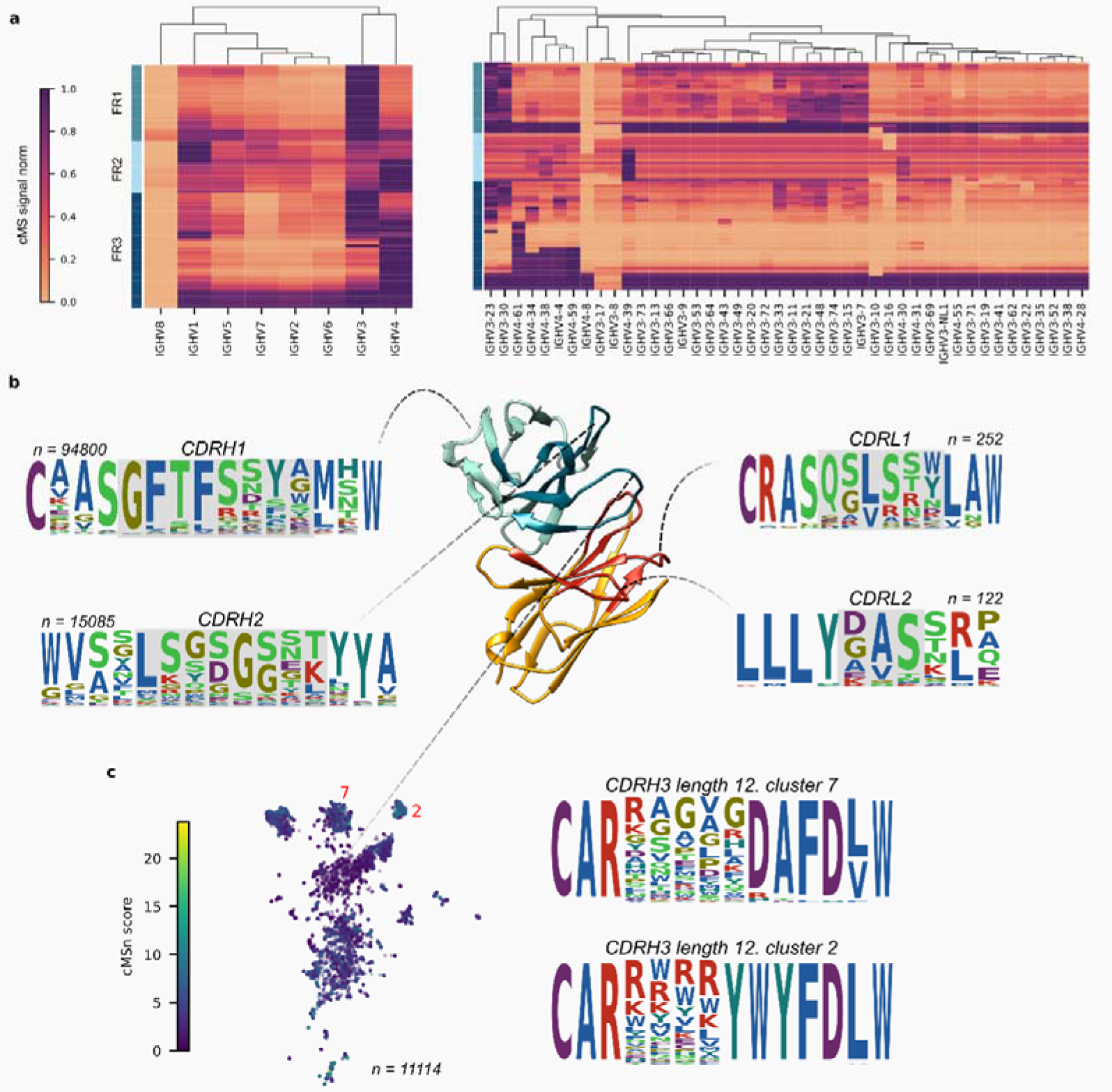
M1-protein binders Fv-population analysis. **a)** Framework cMS signal distribution for all heavy chain groups (left), with a detailed look into the IGHV3 and IGHV4 subgroups (right). **b)** CDR1 and CDR2 feature space distribution via cMSn-based analysis. **c)** exploring 12-mer CDRH3 space. The figure shows the logos for two clusters (2 and 7) in the UMAP-2D embedded space with high cMSn values.

### cMS extract relevant antibody CDR binding features

We then focused on characterizing IgG heavy and light chain CDR populations (Figure 4b). This task is critical to any polyclonal antibody response study, as CDRs are the primary determinant for antigen binding. To achieve this, we walked a built-directed graph between contiguous frameworks, which were constrained by experimentally reported starting and ending 7-mer motifs^52^. For example, for a particular FR1 C-terminal 7-mer, we extracted the experimentally reported CDRH1 lengths and their associated FR2 ending motifs from the IgG database (see *Supplementary information*). We summarized the paths in a CDR-length-specific manner by creating positional MSn-weighted logos from the resulting sequence paths. In addition, Ab25, Ab32, and A49 antibody CDR sequences were used as comparisons, as these antibodies are well-tested and validated M1-binders.

Assembly results suggest that our approach allows us to uncover important positional CDR features (Figure 4b). Heavy and light chains populational data shows conserved feature motifs for CDR1 and CDR2, i.e., length 6 CDRL1 displays a strong signal for the LG motif followed by a polar amino acid, such as N, S, or T; the CDRL2 logo of length three exhibits [A or V]S, a combination of small aliphatic hydrophobic and polar residues, respectively. On the other hand, CDRH1 showed the most conserved features of GFTFS after pooling over 94,800 sequences bearing CDR1 segment of length 8. Finally, the CDRH2 cMS-based logo built from 15,085 sequences highlights L and G in positions 1 and 5, respectively. Interestingly, and even though the experiment selected fragmentation method does not allow us to distinguish between the Leucine (L) and Isoleucine (I) amino acid isomers, we evidenced the presence of the most weighted CDR1 and CDR2 found features in Ab25, Ab32, and Ab49 antibodies. We can see the following tuples:

- CDRL1 (CRAS**QSISSW**LAW, CRAS**QPLSGY**LAW, CRAS**QSVSSY**LAW)
- CDRL2 (LLIY**DAS**SLE, LLIY**NAS**KRA, LLIY**DAS**NRA)
- CDRH1 (CAAS**GFTFSSYA**MHW, CVAS**GFMFNEYY**MSW, CAAS**GFTVSINY**MSW)
- CDRH2 (WVA**LISYDGRNK**YYA, WISF**ISNAGTYT**NYA, WVS**VIYSGGST**YYA).

Regarding the CDR3 region, we identified 416,163 CDRH3s (lengths 8 – 18) and 37,972 CDRL3s (combined IGKVs and IGLVs of lengths 7 – 14). We found a wide diversity of sequences in composition and length compared to the other two CDR regions (Figure 4c). This first finding was expected if we consider the CDR3 region an inherently random gene formation process. In addition, the possibility of finding binders for the different antigen regions was open as the full-length M1 protein was used as bait in the experiment setup. Furthermore, we located up to 35 conserved pairwise features between the aforementioned Ab25, Ab32, and Ab49 CDRH3s when considering 1_2 to 1_8 pairwise interactions (see *CDR3 pairwise amino acid interaction extraction* section). We weighed these pairwise interaction features across all our assembled CDRH3 space. High-weighted interactions reflect the conserved relationship in our population with high MS evidence. For example, the 1_2_RW motif was found in 28,391 sequences (6.8% of CDRH3 space), 1_3_RW (9.3%), 1_3_PK (13.1%), 1_2_RR (27.3%), 1_2_AR (78.4%). These motifs can play a structural or functional role in antibody binding.

Regarding the Ab25 antibody, it bears two high-weight pairwise interactions located in the middle of the CDR3 sequence, suggesting that those are involved in binding. These are 1_3_PK and 1_2_RW motifs with normalized values of 0.82 and 0.74, respectively. Other motifs ranked in the top 5 are located in the beginning or ending part of the sequence, such as 1_2_CA (0.70), 1_3_DW (0.68), and 1_2_AR (0.66). From Ab49, we highlighted the exposure to binding 1_3_RW motif with the normalized value 0.72. For Ab32, the best pairwise interaction motifs are scored below 0.67. These results, along with other CDRs, aligned perfectly with the experimentally reported binding affinity, where Ab25 > Ab49 > Ab32^47^. Finally, some of the above described VH/VL CDR1, CDR2, and CDR3 are also represented in previously reported mouse-derived M1-protein binding antibodies^57^. These results suggest that our approach allowed us to detect CDRH3 sequence patterns of potential importance for targeting the same M1 protein regions.

### cMS integrates structural modeling data for uncovering key antibody-antigen interactions

Our next goal was to sample our detected CDRH3 space to pinpoint potential binders across the included M1 protein domains (A region, B-repeats, Spacer region, and C-repeats). To that end, we tested CDRH3 sequences belonging to different cluster groups as well as spanning a range of lengths and cMS scores. These CDRH3 sequences were grafted onto an IGHV3-21 scaffold, selected for having the highest amount of cMS support (Figure 4a). Following the same principle, a fixed IGKV3-23 scaffold for the light chain was used. After generating heavy–light chain pairs, we employed a modified version of the Sidewinder pipeline’s Structure and Scoring modules^57^. This included structure prediction of the designed Fv and antigen, followed by a targeted molecular docking strategy that scores potential epitopes on a per-residue level based on ensembles of antibody-antigen complexes. Clusters for high-scoring epitopes were then used as the starting point for 500ns Molecular Dynamics (MD) simulations to evaluate the stability of the predicted complexes. Decoys from low-scoring clusters were also selected and submitted to the simulations to assess the rigor of our docking-MD coupled inference evaluation strategy.

The MD simulation results show that our cMS-driven approach identifies stable M1protein binders, over time, across all regions. Given the polar nature of the most studied binding sites, we tracked complex hydrogen bond formation as a proxy for antibody-antigen complex stability. Figure 5a shows candidate antibodies binding M1 protein regions forming inter-protein hydrogen bonds (HB) observed during at least 40% of the MD simulation. Antibody-M1 complexes HB counts indicate that our sampling and modeling strategy has the potential to discover CDRH3s that enable antibody-antigen complex formation and stabilization over time. Furthermore, some CDRH3s targeting the B-repeats or the Spacer region formed up to 11 HB. The simulations also revealed CDRH3s able to bind to two different M1 protein regions, which is consistent with previous findings suggesting dual-Fab cis binding capabilities of the M1 protein^47^. This included members of cluster 7 for CDRH3 of length 12 (Figure 4c), where D104 (a conserved positional cMS amino acid) interacted with the K55 or K85 residues in the A or B-repeat regions, respectively. Another key observations is that our two-step workflow can be utilized as a filter component in an antibody explorative process, which is evidenced in Figure 5a, showing antibody docking results split between the Fv binder and decoy groups.

**Figure 5.**
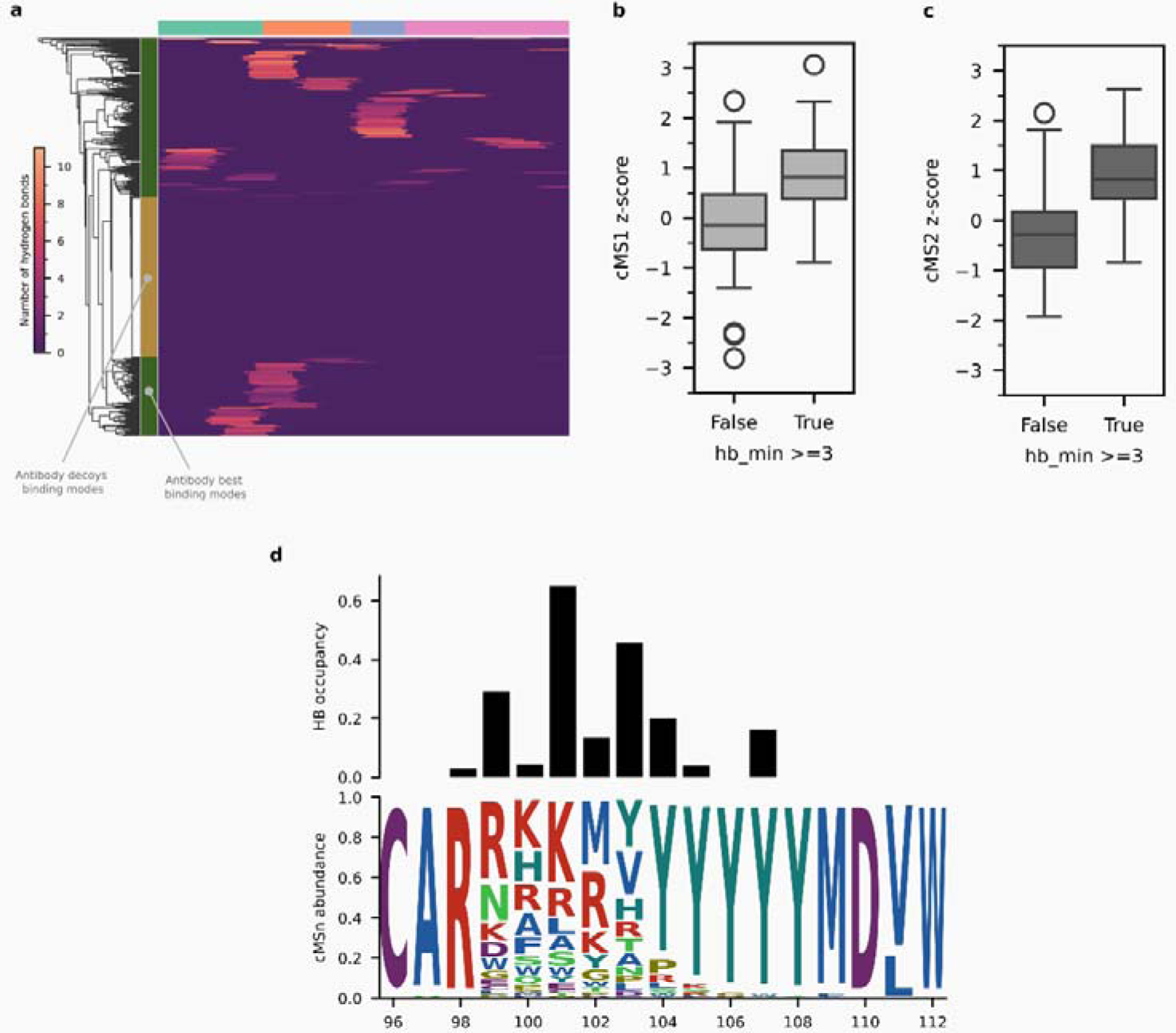
A sampling of the CDRH3 space and M1 protein epitope mapping by MD antibody-antigen complex evaluation **a)** Total number of observed hydrogen bonds (HB) for at least 40% of the MD simulation for all modeled antibody-antigen binding modes (best-selected vs decoys). Columns represent the M1 domain where the antibody binds. CDRH3 cumulative score differences for the groups of antibodies that can form a minimum of three durable HB over time (True group) and those that do not (False group) at the **b)** MS1 and **c)** MS2 levels. **d)** Integrated MS and antibody-antigen modeling data for the sampled CDRH3 of length 15.

It is also worth mentioning that CDRH3 sequences with higher cMS tend to form more HBs than those with low MS evidence. This was observed at both the MS1 (Figure 5b) and MS2 (Figure 5c) levels. Elevated cMS values come from overlapping k-mers with high cumulated MS signals. Given the complexity of our source of antibodies, this indicates that the pulldown experiment, and subsequent computational analysis, captured multiple CDRH3 sequences that share valuable and detectable binding motifs for the M1 protein antigen. This opens a new venue for exploratory antibody studies that shifts the focus away from complete characterization of the B-cell repertoires, or precise *de novo* MS monoclonal antibody assembly, towards feature optimized antibody design based on biologically constrained systems.

Finally, it is possible to integrate all the acquired data layers from our system of interest and uncover potential position-wise information. For example, the combined results for the sampled CDRH3 sequences of length 15 (summarized in Figure 5d), revealed that position 101 was the most persistent paratope residue to form HBs with the antigen at this length. This observation is backed up by high cMS positional abundance values for both K and R amino acids. In other words, having either of these two amino acids in that position of the CDRH3 sequence makes the antibody able to form a persistent hydrogen bond with the M1 antigen. In addition, other polar amino acids with high cMS abundance, such as R99 and Y103, were also identified in other CDRH3 length clusters with high cMS (see cluster 2 in Figure 4c).

Our comprehensive analysis demonstrates a remarkable consistency between the *de novo* MS peptide sequencing annotation method, the cumulative MS signal analysis, and established genetic and experimental findings. The agreement between the high cMS values, the identified VH and VL subgroups, and the experimentally validated CDR features underscores the robustness of our approach. Furthermore, the ability to pinpoint conserved CDRH3 motifs and predict stable antibody-antigen interactions through docking and MD simulations highlights the method’s potential for exploring complex polyclonal antibody responses. The integration of these data layers offers promising avenues for uncovering position-wise information and developing novel antibody selection strategies. Exploring related use cases, such as BCR epitope prediction based on sequence alone, could also be of interest. State-of-the-art protein-protein interaction predictors that rely on multiple-sequence alignments, like AlphaFold-multimer^21^, fail to reliably predict antibody-antigen complexes due to the lack of coevolutionary signal^22^. Our results suggest that there might be a paratope-epitope sequence feature space to mine, similar to previously published structural feature mining ^58^, that could be used to develop novel prediction models. The insights gained from this study, particularly the correlation between cMS values and hydrogen bond formation, suggest that leveraging cumulative MS signal analysis can significantly enhance our understanding of antibody-antigen interactions and facilitate the discovery of potent therapeutic antibodies.

## Conclusions

We present a novel approach for successfully annotating *de novo* sequenced MS peptides, evaluated and validated comprehensively across diverse sample types and experimental conditions. This methodology, transitioning from isolated spectrum-matching and string-based to MS signal-based analysis, allowed us to effectively characterize complex protein systems, particularly a polyclonal antibody mixture of *S. pyogenes* M1 protein binders. By leveraging cumulative MS signal (cMS) values, we identified predominant VH and VL subgroups, aligning with established genetic studies and experimental observations. Furthermore, our approach revealed conserved CDR features, including length-specific motifs within CDR1 and CDR2, and uncovered a wide diversity of CDR3 sequences, reflecting the inherent randomness of this region. Notably, we pinpointed CDR3 motifs with potential binding capabilities, correlating high cMS values with stable antibody-antigen interactions, as evidenced by molecular dynamics simulations. This consistency across multiple data layers, from subgroup identification to binding prediction, underscores the robustness of our method. Moving forward, integrating these insights offers promising avenues for refining antibody selection strategies and gaining deeper positional knowledge of antibody-antigen interactions, paving the way for new targeted therapeutic development.

## Material and methods

### Computational details

#### Cumulative MS scoring system

Let’s consider a set of spectra *s*, defined as s = {s_1_,s_2_,…, s_m_}. *S* represent the combination of all spectra acquired using different proteases and experimental set ups. In addition, we have a series of candidate peptides *c* given by *c_m_ = {c_1n_, c_2n_, …, c_mn_}* for every spectrum *s_m_*. The set *c* come from the combination of different spectrum-matching engines, like database-based or *de novo* MS peptide sequencing tools. We define our novel cumulative MS score (cMS) for every *c_mn_* as:

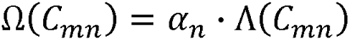

Here, we call Λ the isolated score and measure the spectrum-matching between the peptide *c_n_* and the spectrum *s_m_*. The factor *a_n_* is the context weight of the peptide *c_n_* when cumulating all the considered MS signal, represented by Γ. Peptides belonging to the target proteins end up with higher context values respect to decoys. In principle, the goal of Ω is to be higher for a true-peptide with respect to a set of decoy-peptides for any spectrum or Ω(*c_true_*) > Ω(*C_decoys_*). For calculating every *a_n_*, we initially create a vector of length *L_n_* for each *c_n_* and fill it with MS evidence, given by Γ*_nm_* = [Γ*_nm_*^1^, Γ*_nm_*^2^,…,Γ*^L^_nm_*]. All positions Γ*_nm_* are a cumulated value from the fragment ions information. Also, for fragment ions *a, b* or *c* data are store in the same position *h* as their numbering *t*, while for *x,y,z* we applied *h = Ln - t*. Then we generate *N* fragments of size *K* given by c1, c2, …, cy] and slice Γ*_nm_* from 1 to L, k + 1, so every fragment is a new vector fi(c) = Γ*_nm_*[i:i+k-1]=[Γ*_nm_^i^*,Γ*_nm_^i+1^*…, Γ*_nm_^i+k^*]. All the stored MS signal in every *f(c*) is then pooled element-wise across all *c_mn_* in *s* to create the new vector of F(c) = Σ^m^, Σ^n^ f*_uv_*(c). We use the newly created *F(c)* cumulated MS vectors to calculate *a_n_* for every observed candidate peptide *c_n_*. We create a vector *Q_n_* of length *L_n_* by aligning all *F* composing *C_n_*. For every position *i* in *Q_n_*, we align a set of *F_p_^q^* given by the formula:

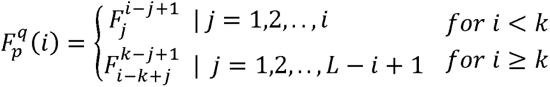

Here, *p* goes from 1 to L – k +1, while 6, the component in *F*, goes from 1 to *k*. Then, for every *i* in *Q_n_*, we apply a fragment operator Φ to produce a context vector Χ(*C_n_*) = [Φ_1_,Φ_2_, …, Φ_L_] for *C_n_* of length *L_n_*. Finally, the context weight is calculated by applying a function on *X(C_n_)*, given by an = (*X*(*C_n_*)). We can make this integrative process iteratively and keep the best candidates according to their Ω(C*_nm_*) values. This new math framework rely on the idea that every peptide has a unique *a* value. From cumulating all the MS signal, true peptides end up with higher *a* values respect to the decoy peptides. Also, context weight values play a critical role in those situations where, due to the lack of fragment ions, the true and decoy peptides share the same isolated score.

In the present work we tested several simple Λ functions by using MS/MS features. MS H_2_O, b-H_2_O, y-H_2_O, a-NH_3_, b-NH_3_ and y-NH_3._ The isolated score was always the Γ_nm_ vector signal came from considering 12 fragment ions, namely: a+1, b+1, y+1, a+2, b+2, y+2, a-mean value of the selected MS signal. For example, in the function 1 we fill the vector Γ*_nm_* adding is a exponential weighted value of Δm, the absolute difference between the observed m/z value respect to the theoretical one, given by S1 = e^-w|Δm |^. We derived S5 for the case of w = 0 in S1. Another MS/MS feature considered was the intensity of the fragment ions. For that we created 4 using the normalized intensity as signal. Furthermore, in a effort to integrate both m/z and intensity features, we fill the vector in 4 with the sum of the amount of observed fragment ions and their intensity. Furthermore, in sS4b, we weighted the contribution of the fragment ions. Here, we tested doubling the contribution of the *a, b*, and *y* single-charged fragment ions, respect to the other fragment ions. Finally, in E-score, the positional signal was normalized respect to the expected number of fragment ions in each position.

On other hand, we used a fragment size *k* of 5 for all our *in silico* experiments. The minimum value of the aligned *F(C)* vectors was considered as operator Φ. In addition, *a* value was the mean value of the resulting vector *Q*. The selection of Φ and *X* aforementioned operators guaranteed that high *a* values come from peptides with many kept top 100, 50, 20, 10, 8, 5, 4, 3, 2, 1 ranked according to Ω, respectively. An Snakemake contiguous highly MS signaled fragments. Finally, we considered 10 iterations where we workflow^59^ with the implementation of the cMS concept can be freely accessed through GitHub https://github.com/cguetot/cms.git.

#### MSn score calculation

Because of possible assembled paths come from overlapping the final annotated peptides, we used the final *Q_n_*, vectors to make downstream population analysis (Figure 4). We Because of possible assembled paths come from overlapping the final annotated peptides, distinguish MS1, MS2 and MSn type of *Q_n_*, vectors created from *F(C)* vectors from the final annotated peptides. MS2 are created using the 12 considered fragment ion information. For fragments of size *k* and assigned the same signal for all components of the generated MS1, we extract the log10 value of the MS1 intensity for all peptides. Then, we created fragment vectors. Finally, we kept the maximum value for every fragment. Furthermore, we multiplied element-wise the normalized values of MS1 and MS2 vectors. *Q_n_* resulting created MSn in a effort to combined MS1 and MS2 into one unified value. For that, we vectors with consecutive high MSn values represent segment with high overall MS evidence. In other words, those assembled sequences have a lot of experimental support from our *de novo* sequencing process. On other hand, we created logos for CDR regions from the assembled populations using the position-wise proportion of the cumulated signal for all amino acids found. Unlike the conventional string-based analysis, high abundance value reflects high MS evidence from the identified population (Figure 4b).

#### CDR3 pairwise amino acid interaction extraction

For each CDRH3 amino acid sequence, pairwise interactions were extracted based on the relative positions of amino acids within the sequence. Given a sequence *S = s_1_ s_2_ … s_n_*, where *s_i_* represents the amino acid at position *i*, all possible pairs of amino acids(*s_i_,s_j_*) were generated, where 1 < i < j < n. The interaction was then defined as a tuple (1, j – i + 1, *s_i_,s_j_*). For example, some extracted pairwise interaction for the sequence CDRSEQPEPTIDE are 1_1_CD, 1_2_ID, 1_3_QE, 1_4_RQ, 1_6_ET, etc. We then assigned an *interaction weight* by taking the minimum value from the quantities in the positions *i* and *j* of the sequence MSn *Q_n_*, vector. Finally, for each unique pairwise interaction, the highest calculated weight was retrained. These high-weight interactions were then analyzed to identify potential motifs related to CDR binding. Specifically, interactions with high weights located towards the center of the CDR3 loop were considered potential binding motifs. Conversely, high-weight interactions occurring near the N-terminal or C-terminal regions of the CDR3 sequences were interpreted as potentially reflecting structural constraints or requirements rather than direct binding interactions.

#### Peptide candidates list generation

MS/MS spectra amino acid sequences were created using database and *de novo* spectrum-matching searches. DDA raw files were initially converted to Mascot generic format (MGF) by ThermoRawFileParser software^60^. On the other hand, employed DIA-Umpire^61^ to convert raw DIA files into searchable MGF format. Comet engine (version 2023.01.02^62^) was used for searching against the target databases. For the Herceptin experiment (Figure 2), we created a combined database with 54256 entries by appending the monoclonal antibody heavy and light chain sequences along with five proteomes with taxonomy identifiers 10090, 559292, 7227, 83333 and 9606. The database expanded peptide list creation was done in a two-steps process: Initially, we made independent searches with every amino acid as C-term ranging the length of 5 to 40. Regarding modifications, optional Met Oxidation (UniMod: 35). For all alkylated samples, the fixed Cys carbamidomethylation (UniMod: 4) modification was considered. We then create a new fasta file with the candidates found and made a intact peptide search adding additional modifications such as UniMod:27 for Glu, UniMod:35 for Met, Tyr, Lys and Arg amino acid. Also, N-term UniMod:1, UniMod:5, and UniMod:34 modifications. On other hand, DeepNovoV2^24^ and CasaNovo^33^ deep-learning-based *de novo* MS peptide sequencing tools were used. For the first one, five SEMs (trypsin-SEM, chymotrypsin-SEM, elastase-SEM, gluc-SEM and pepsin-SEM) and two MEMs previously trained were used^42^. The DeepNovoV2 code was modified to expand candidate peptides list for each spectrum. For CasaNovo, all searches were performed using the default model and parameters. For each spectrum, top 20 scored results were considered.

#### Antibody structural modeling

For the Herceptin full-length model, the Fc and Fab domains were de novo modeled separately by AlphaFold2^63,64^, using MMseqs2^65^ to generate the multiple sequence alignment and homo-oligomer state of 1:1. For each selected model, the sidechains and the disulfide bridges were adjusted and relaxed using the Rosetta relax protocol^66^. The loops in the hinge region were then re-modeled and characterized using DaReUS-Loop web server^67^. Finally, the full-length structure was relaxed, and all disulfide bridges (specifically in the hinge region) were adjusted using the Rosetta relax protocol. To showcase the benefit of adding more data from different *de novo* MS peptide sequencing models and tools, we calculated the *confident positional score* (CS^42^) for all *de novo* peptide cMS results coming from SEMs, CasaNovo, MEMs and the combination of all of them (NOVOs). For every position *i* in the amino acid sequence, *CS_i_* = log_2_ (f_i_ +1) is defined as logarithmic transform of the number of *de novo* sequenced matching peptides found in the position *i* of the target protein sequences (*f_i_*). Higher consecutive CS values represent regions with more evidence in the *de novo* protein sequencing process, being especially important for MAbs HC and LC variable regions^42^. Visualization of the Herceptin monoclonal antibody with the confident score layout was done through USCF Chimera^68^ software (Figure 2g).

#### Sidewinder-based populational docking and scoring

To identify antibody epitopes, we employed a strategy similar to the one implemented in the Sidewinder pipeline^57^, excluding the cross-linking mass spectrometry (Data) module. Specifically, we used MEGADOCK^69^ (v4.1.0) to perform molecular docking with IgFold^70^ (v0.4.0) predicted Antibody Fv’s and a AlphaFold3^71^ (accessed 11.11.2024) predicted truncated M1 protein structure. This truncated version included residues P18–E307, and two glycine were added at both the N and C-terminus of both chains to reduce the molecular docking bias towards the termini. To limit the predicted interfaces to the Fv CDR residues, we annotated the Fv sequences using the IMGT numbering scheme^72^ and masked framework residues before running MEGADOCK. In our population approach, we calculated a weight *W(a)* for every position *a*, following the formula *W(a)* ∑_pep_ H(a,p), where framework residues before running MEGADOCK. In our population approach, we calculated *W(a)*. Here *p* is the set of the top N best binding poses according to CDRs. The term *d(a,c)*, denotes the Euclidean distance between the alpha-carbon of the the MEGADOCK scoring system, and CDR is the set of amino acid *a* positions in the antibody antibody-antigen interaction for the pose *p* according to ZRANK2^73^. The score *H(a,p)* antigen amino acid and CDR amino acid, while represents the rescored value of the calculates the contribution of antigen position *a* in pose *p*, and *W(a)* is the summation of these contributions across all poses, effectively highlighting antigen positions that frequently and closely interact with the antibody CDRs across the top N binding results. For each Fv-antigen pair, 2000 conformations were predicted, and the top 200 decoys (the population) were scored using the aforementioned modified version of the Sidewinder Scoring module.

#### Molecular Dynamics simulations

Given the long coiled-coil structure of the M1 protein, we designed a fragmented version for our MD simulations pipeline to avoid using the full-length M1. We preserved the IgG variable domain and retained only M1 regions within 10 Å of IgG, removing the rest. If this threshold caused partial fragmentation or disrupted the coiled-coil structure, we increased it accordingly. As a result, our MD simulations used partial M1 domains interacting with IgG variable domains. For every system, we performed three replicates of 500 ns MD simulations starting from the input structure, explained in the previous section. MD simulations were carried out with the GROMACS 2024.2^74^ using the CHARMM36m force field parameter set: (i) Na+, Cl-counter-ions were added to reproduce physiological salt concentration (150 mM solution of sodium chloride), (ii) the solute was hydrated with a dodecahedron box of explicit TIP3P water molecules with a buffering distance of up to 12 Å, and (iii) hydrogen atoms were added and the environment of the histidine was checked using the Reduce software^75^. For each system, the energy minimization was performed by the steepest descent algorithm for 5000 steps, to minimize any steric overlap between system components. This was followed by an equilibration simulation in an NPT ensemble at 310 K, allowing the lipid and solvent components to relax around the restrained protein. All the protein and lipid non-hydrogen atoms were harmonically restrained, with the constraints gradually reduced in 6 distinct steps with a total of 0.375 ns. During the system equilibration steps, the pressure was maintained at 1 bar with Berendsen thermostat and barostat^76^, respectively. For every system, three replicates of 500 ns, with different initial velocities were performed in the NPT ensemble using the time step of 2.0 fs. The temperature was kept constant at 310 K using V-rescale thermostat^77^ and a constant pressure of 1 atm was maintained with C-rescale barostat^78^. Isotropic pressure treatment was applied to the system. The bonds involving hydrogen atoms were constrained using the LINCS algorithm^79^, while the electrostatic interactions were calculated using Particle Mesh Ewald method^80^, and the coordinates of the system were written every 100 ps.

#### Antibody-Antigen hydrogen bonds detection

The hydrogen bonds (HB) were detected using the HBPLUS algorithm^81^. HBs are detected between donor (D) and acceptor (A) atoms that satisfy the following geometric criteria: (i) maximum distances of 3.9 Å for D-A and 2.5 Å for H-A, (ii) minimum value of 90° for D-H-A, H-AAA and D-A-AA angles, where AA is the acceptor antecedent. For a given HB between residues i and j, interaction strength is computed as the percentage of conformations in which the HB is formed between any atoms of the same pair of residues (i and j).

### Experimental details

#### Multi-enzyme Digestion of Monoclonal Antibody Avastin

A total of 500 µg of the monoclonal antibody Avastin (Roche) was processed by standard in-solution reduction and alkylation prior to different protease digestion. The antibody was first diluted in 8 M urea (Sigma-Aldrich) 100 mM ammonium bicarbonate (Sigma-Aldrich) buffer and then incubated with 5 mM tris(2-carboxyethyl)phosphine (TCEP; Thermo Fisher Scientific) for 1-hour reduction at 37 °C and 300 rpm in a ThermoMixer (Eppendorf). This was followed by alkylation with 10 mM iodoacetamide (IAA; Sigma-Aldrich) in the dark for 30 minutes at room temperature. The sample was subsequently aliquoted for multi-enzyme, multi-time-interval digestion.

Digestions were performed using the following conditions: for trypsin (Promega), chymotrypsin (Promega), and alpha-lytic protease (Sigma-Aldrich), 100 mM ammonium bicarbonate was used as the digestion buffer; for elastase (Promega), 50 mM Tris, pH 8.0; for Glu-c (Promega), 100 mM Tris (Sigma-Aldrich) pH 8.0; and for pepsin (Promega), 0.04 N HCl (Merck) at pH 1.5. Three digestion intervals (4 hours, 9 hours, and overnight >14 hours) were applied for each enzyme, except for pepsin, which instead underwent digestion intervals of 15, 30, and 60 minutes. The protease-to-protein ratio was set at 1:20 (w/w) for all enzymes, except for pepsin, for which a ratio of 1:6 was used. Digestion was terminated by acidification with formic acid (Fisher Scientific) or heating to 95 °C. The resulting peptides were purified using a C18 spin column (Thermo Fisher Scientific), concentrated and dried using a SpeedVac (Eppendorf), and reconstituted in buffer A (2% acetonitrile, 0.2% formic acid; Thermo Fisher Scientific) for subsequent mass spectrometry analysis. Approximately 1 µg of the mAb digest, as determined by a NanoDrop spectrophotometer (DeNovix), was loaded onto an Ultimate 3000 UPLC system (Thermo Fisher Scientific) interfaced with a Q Exactive HF-X hybrid quadrupole-Orbitrap mass spectrometer (Thermo Fisher Scientific). Peptides were first concentrated on a precolumn (PepMap100 C18 3 μm; 75 μm × 2 cm; Thermo Fisher Scientific) and then separated on an EASY-Spray column (ES903, maintained at 45 °C; Thermo Fisher Scientific) according to the manufacturer’s recommendations. Mobile phases comprised solvent A (0.1% formic acid) and solvent B (0.1% formic acid, 80% acetonitrile). A linear gradient from 3% to 38% solvent B was applied over 120 minutes at a constant flow rate of 350 nl/min.

#### M1-specific IgG pulldowns

M1-specific IgG was purified from IVIG as previously reported^50^. Roughly, 20 µg of recombinantly expressed M1 was immobilized on pre-equilibrated AssayMAP Streptavidin columns (Agilent Technologies). Columns were washed with PBS and then IVIG (100 µg) was applied, followed by extensive PBS wash. Elution was done using 100 µl of 0.1 M glycine (pH = 2) and the final pH was neutralized with 20 µl of 1 M Tris. Samples were reduced with 5 mM Tris(2-carboxyethyl) phosphine hydrochloride (TCEP), alkylated with 10 mM iodoacetamide, and digested with trypsin (1:100) at 37 °C for 18 h, and the digestion was stopped using 20% TFA (Sigma) to pH 2–3. Peptide clean-up was performed using AssayMAP C18 columns (Agilent Technologies) per the manufacturer’s protocol. Samples were dried using vacuum concentrator (Eppendorf) and resuspended in 20 µl 0.1% formic acid (FA, Fisher Chemical) followed by a brief sonication for 5 min before analyzing on a Q Exactive HF-X mass spectrometer (Thermo Scientific). Following the aforementioned multi-time procedure, digestions were performed using trypsin, chymotrypsin, elastase, and pepsin.

#### Mass Spectrometry Analysis

Data acquisition was performed using either a data-dependent acquisition (DDA) or a data-independent acquisition (DIA) method. In the DDA method, an initial MS1 scan (350-1650 m/z, resolution of 60,000, AGC target of 3e6, maximum injection time of 50 ms) was followed by the top 20 MS2 scans (resolution of 15,000, AGC target of 1e5, injection time of 25 ms) with a stepped normalized collision energy (NCE) of 20, 26, and 35. Charge states of 6 to 8, and those above were excluded, except for samples digested with trypsin enzyme, in which singly charged ions were also excluded. For the DIA method, a shorter gradient from 3% to 38% solvent B was applied over 90 minutes at a constant flow rate of 350 nl/min. An MS1 scan (390-1210 m/z, resolution of 120,000, AGC target of 3e6, maximum injection time of 60 ms) was acquired, followed by MS2 scans across 44 variable isolation windows (resolution of 30,000, AGC target of 3e6, auto-mode injection time) with a stepped NCE of 20, 26, and 35. The performance of the LC-MS system was monitored by analyzing a yeast protein extract digest (Promega). Datasets were acquired in a consecutive run.

## Acknowledgment

This research was supported by the Swedish Research Council (Vetenskapsrådet (VR); VR-2020-02419 and VR-2022-05264), the Knut and Alice Wallenberg (KAW) Foundation (KAW 2016.0023), and the Alfred Österlunds Foundation. J.M. is a Wallenberg Academy fellow (KAW 2017.0271) and is also funded by the Swedish Research Council (VR; 2023-02107), the Wallenberg Foundation (KAW 2019.0353), and Alfred Österlunds Foundation.

